# Single molecule studies of the bacterial curli protein CsgA reveal a structurally dynamic monomeric structure

**DOI:** 10.1101/2025.09.08.674806

**Authors:** Daniel A.N. Foster, Kirill Kudriavtsev, Calum Byrne, Simcha Srebnik, Derek R. Dee

**Affiliations:** Food, Nutrition and Health, Faculty of Land and Food Systems, The University of British Columbia, 2205 East Mall, Vancouver, BC, V6T 1Z4, Canada; Chemical and Biological Engineering, The University of British Columbia, 2360 East Mall, Vancouver, BC, V6T 1Z3, Canada

## Abstract

*E. coli* curli, an archetypical bacterial amyloid, is a critical biofilm scaffold comprised mainly of the CsgA protein. CsgA is highly aggregation prone, confounding high-resolution studies of its structure and amyloid formation mechanism. Ensemble studies show that monomeric CsgA is intrinsically disordered, while structure predictions show a well-folded β-solenoid. Here, single-molecule force spectroscopy with optical tweezers is used to define the conformational dynamics in single CsgA molecules. CsgA monomer shows unfolding/refolding consistent with fully folded, partially folded and collapsed molten globule conformations, indicating that CsgA exists in metastable folded states in equilibrium with disordered states. These folded states are likely highly aggregation prone and thus have avoided characterization at the ensemble level. Steered molecular dynamics simulations reveal unfolding patterns consistent with the experimental data. These studies help us better understand curli amyloid formation that is important both as a novel antimicrobial drug target and engineered biomaterial.

## INTRODUCTION

Bacteria within biofilms are more resistant to antibiotics and are a major health concern^1^. Curli fibrils, a functional amyloid, are a key biofilm component and novel drug target^2^. Curli, as a genetically programmable scaffold, is also of interest as a novel biomaterial^3^. *E. coli* curli is the best studied functional amyloid system, involving several chaperones encoded by the curli-specific genes (csg) operon that direct the assembly of fibrils comprised mainly of CsgA. Inside the periplasm, CsgA aggregation is inhibited by the chaperone proteins CsgC and CsgH^4^. The secretion of CsgA to the outer membrane is controlled by CsgG, CsgE and CsgF^5^, where CsgB nucleates CsgA fibril formation on the cell surface^6^. CsgF is anchored to the outer membrane and may in turn anchor CsgB^7^. While these biological pathways are known, many of the mechanistic details are not^6,8,9^, such as the dynamic structure of CsgA, how CsgB nucleates CsgA amyloid formation, and how CsgC, CsgE and CsgH inhibit CsgA fibrillation^8,10^.

The CsgA sequence comprises a signal peptide (res. 1–20) that is cleaved, a CsgG-binding N-terminal domain (N22)^11^, and an amyloid core region composed of five imperfect repeats (R_i_) each 19-23 aa long (**Fig. 1a**). R1 and R5 govern CsgA responsiveness to CsgB nucleation and self-seeding by CsgA fibers and can aggregate in isolation, being more aggregation prone than R2-R4 that are inhibited by several Asp and Gly “gatekeeper” residues^12^.

**Figure 1.**
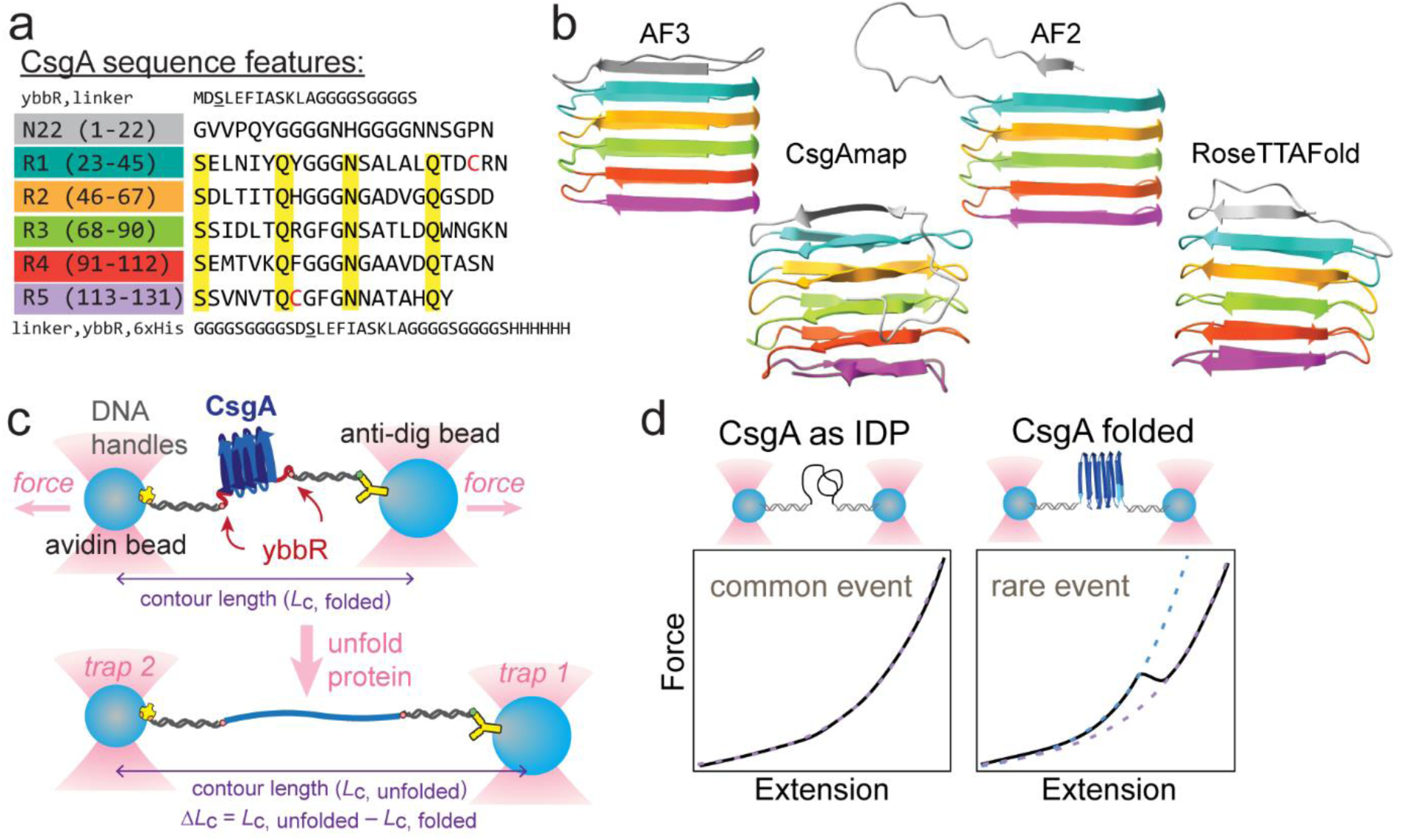
CsgA sequence features, predicted structures and single molecule force spectroscopy. (**a**) Sequence features of CsgAcc engineered for single molecule study. Residue numbering begins with the mature N-terminus (signal sequence omitted). (**b**) Predicted CsgA structures: AF2, AF3, RoseTTAFold and DeBenedictis 2019. (**c**) Dual-beam optical tweezers experiment with the protein tethered between two trapped beads using dsDNA handles. Moving the beads apart unfolds the protein and the change in contour length (Δ*L_c_*) is determined. **(d)** Boundary conditions for what can be expected when measuring CsgA: no transitions (protein unstructured through the entire measurement) or discrete transition (protein is folded at low force, then unfolds giving a transition and continues to be stretched out to high force).

Understanding protein folding requires knowing the folded structure and resolving the folding pathway; the former can now generally be predicted with high confidence while the later remains a challenge^13^. For CsgA, various computational models predict the monomer to form a β-solenoid structure^14–17^ (**Fig. 1b**). This structure is consistent with cryoEM data of CsgA amyloid fibrils^18^, suggesting that folded CsgA monomers, if not CsgA dimers^19^, may constitute the minimal amyloid unit. From ensemble studies, monomeric CsgA is found to be intrinsically disordered^6,9,20,21^ and rapidly fibrillates into a β-sheet rich amyloid. The first steps of nuclei formation are hypothesized to include either transient folding of two CsgA monomers that then bind together to form the amyloid nucleus, or co-operative binding and folding of two CsgA monomers to form structured dimers^9^. The folding and amyloid-forming pathway of CsgA remains to be resolved.

Ensemble structural methods report the average features for billions of unsynchronized molecules, limiting the resolution of molecular-level details in fibril formation due to the inherent heterogeneity of aggregation^22,23^. Experiments on single molecules avoid ensemble averaging, allowing otherwise invisible features such as molecular distributions, specific pathways, and rare or transient structures to be observed^23^. Optical tweezers (OT) are well-suited for single-molecule force spectroscopy (SMFS) studies (wherein individual biomolecules are unfolded and refolded under force) due to their high resolution in force (sub-pN), space (~0.1 nm), and time (<1ms)^23^. OT have been used to resolve native folding and misfolding conformational energy landscapes^24^ and to uncover critical early steps in the aggregation of disease-related proteins such as neuronal calcium sensor-1 (linked to schizophrenia and bipolar disorder)^25^, α-synuclein (Parkinson’s)^26^, prion protein (transmissible spongiform encephalopathies)^27,28^, and superoxide dismutase (amyotrophic lateral sclerosis)^29^. Avoiding ensemble averaging is particularly useful to resolve the conformational dynamics of intrinsically disordered proteins (IDP’s), and these have been studied at the single molecule level using fluorescence, nanopores, and SMFS^30^. SMFS with OT has been used extensively to study Aβ_1-42_ and α-synuclein^24^, two IDP’s that form pathological amyloid. SMFS, using magnetic tweezers, was used to examine the mechanical properties of curli nanofibers^31^ and to probe unfolding of a single CsgA molecule, indicating stepwise unfolding consistent with a completely structured molecule^32^.

Here we use SMFS with OT to probe the structure of monomeric CsgA. In a typical OT setup, a biomolecule of interest is tethered between two DNA handles attached to μm-sized beads trapped by tightly focused lasers (**Fig. 1c**). Force is then controlled via changing the distance between beads, recording force extension curves (FECs), and the response of biomolecules is used to reconstruct detailed folding/unfolding pathways^33^. CsgA was expected to exist between two boundary conditions (**Fig. 1d**): as an unstructured IDP, showing featureless FECs similar as monomeric α-synuclein^26,34^, or showing discrete transitions (a “rip” in the FECs) consistent with the predicted folds (**Fig. 1b**) but never before observed experimentally. We resolve complex conformational dynamics that would otherwise be hidden by ensemble averaging and elucidate (un)folding pathways that may represent the earliest steps of CsgA amyloid assembly.

## RESULTS

To restrain CsgA aggregation before single molecule tethering, we developed an assay to trigger native folding of CsgA under controlled conditions in a microfluidic flow cell. This work exploits an engineered disulfide trap in the protein (CsgA-A43C/V120C, termed “CsgA_CC_”)^35^, where fibrillation becomes possible only after reduction of the cystine. Introducing a disulfide bond between the highly amyloid-prone R1 and R5 regions of CsgA (A43, V120, red **Fig. 1a**) facilitated sample preparation (recombinant expression/purification, bioconjugation with handles, and isolating single molecules) by limiting premature aggregation. At the ensemble level, fibrillation is induced by adding reducing agent and, otherwise, the mutations do not affect CsgA fibrils^18,35,36^. DNA handle attachment proceeded using engineered ybbR tags with CoA-modified oligonucleotides and the 4’-phosphopantetheinyl transferase Sfp chemistry^37^. The ybbR-CsgA_CC_-ybbR construct was expressed and shown to form amyloid only upon adding TCEP (**Fig. S1**). After attaching dsDNA handles, the CsgA construct was introduced into the microfluidic flow cell and tethered between two polystyrene beads. CsgA constructs were initially screened for a single tether in running buffer without reducing agent. The data was fit offline to a worm-like chain (WLC)^38^ to confirm that it was a single tether (**Fig. S2**). The FEC data were fit using a polymer physics model (Marco-Siggia WLC^39^) to describe the elastic compliance of DNA and protein in series, permitting the measurement of protein Δ*L_c_* upon unfolding or refolding. Before reduction, with the internal disulfide intact, FECs were typically smooth with no apparent transitions. After the addition of a reducing agent (**Fig. S2b**), the FECs showed transitions and a total contour length consistent with the length of the dsDNA plus reduced protein construct. Seldom, transitions were observed in the disulfide-linked CsgA (**Fig. S2c**), with a total contour length change (Δ*L_c,_*_tot_) consistent with R1 and R5 forming a structure together. From there we measured the individual unfolding and folding trajectories of single molecules of CsgA using OT. We analysed data from 15 CsgA molecules with over 800 pulls overall, examining the effects of pulling velocities (50, 100, 200, 300 and 500 nm/s) and refolding times at low force (200 ms to several seconds) on the unfolding of CsgA.

### CsgA shows discrete and non-discrete transitions

The unfolding of CsgA showed three general behaviours (**Fig. 2**): FECs with only a non-discrete “shoulder”, FECs with a shoulder lead-in followed by ≥1 discrete transition, and FECs with only discrete transitions. The total Δ*L_c_* was similar for all three behaviours (~44 nm, **Fig. 2a**) but comprised of varying amounts of shoulder and discrete transitions. Within the two discrete sets, FECs showed a variable number of transitions, with some showing one, two or three discrete unfolding transitions (**Fig. 2b-d**), with most showing just a single transition. Overall, 67% of FECs showed at least one discrete event and 33% showed no discrete events (shoulder only) (**Fig. 2e**). Thus, all FECs showed some type of transition and the featureless (monotonic) extension curves expected for a random coil (**Fig. 1d**) were not observed here.

**Figure 2.**
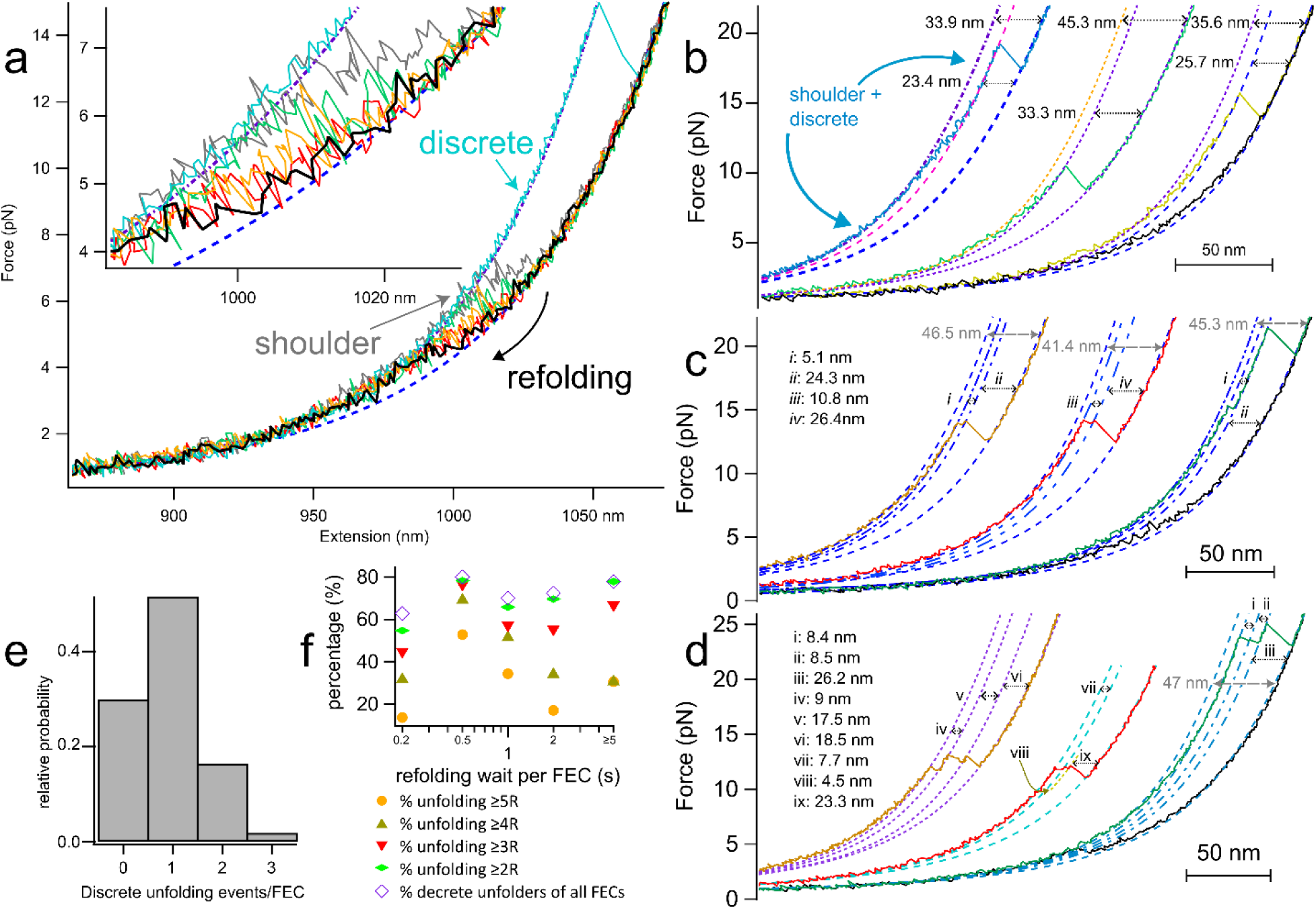
Unfolding single molecules of CsgA. (**a**) FECs showed three general behaviours: non-discrete unfolding (also refolding) “shoulder” transitions, a shoulder followed by discrete transitions, and (rarely) discrete only. (**b-d**) Examples of FECs with varying numbers of discrete unfolding events. (**e**) Relative probability of FECs showing a shoulder only *vs* one, two or three discrete transitions. (**f**) Percentage of FECs with a discrete transition of any Δ*L_c_*__tot_ and with a Δ*L_c__*_tot_ consistent with ≥ 2, 3, 4 or 5 folded CsgA repeats, as a function of refolding time at low force. Dashed lines in (**a-d**) show WLC fits.

In example FECs with a single discrete rip fit with a WLC (**Fig. 2b**), in some cases the data diverges from the WLC fit at low force owing to the shoulder transitions (blue), while in others (yellow, green) no shoulder is evident. Fitting the entire folded arm (WLC fit leading up to the shoulder) gives a total Δ*L_c_* of ~44 nm indicating involvement of the entire protein, including the N22 domain (**Fig. S3, S7**). In FECs with two (**Fig. 2c**) and three (**Fig. 2d**) discrete unfolding steps, in both cases the discrete unfolding events were (nearly) always preceded by a low-force shoulder transition. Sequential transitions typically occurred within a narrow range of forces (**Fig. S4**), a consequence of subsequent transitions occurring shortly after the first ones; less often, the final transitions occurred at much higher forces.

The refolding waiting time (time spent at low force between successive pulls) had no measurable effect on the probability of FECs showing larger transitions (*e.g*. those corresponding to ≥3 or ≥4 folded repeats) (**Fig. 2f**). This suggests these structures reach equilibrium faster than our observation window and that they are metastable relative to each other and to the rapidly interconverting states that give rise to the shoulder transitions (discussed below).

The unfolding forces and contour length changes for individual discrete transitions (**Fig. 3a**) covered a broad range, from 5 pN to 35 pN and 4 nm to 46 nm, respectively. The total discrete Δ*L_c_* (sum of 1-3 transitions) similarly spanned from 4 nm to 46 nm with several peaks evident in the distribution (**Fig. 3b**). These peaks correspond to the expected Δ*L_c_* for integer and half-integer amounts of CsgA repeat structure (**Fig. S3**), ranging from one folded repeat (1R) all the way to 5R, 5R + half of N22, and 5R + N22. Clustering of contour length changes at 23, 30 and 37 nm could be attributed to 3, 4 and 5 repeats, respectively, as shown (**Fig. 3b**).

**Figure 3.**
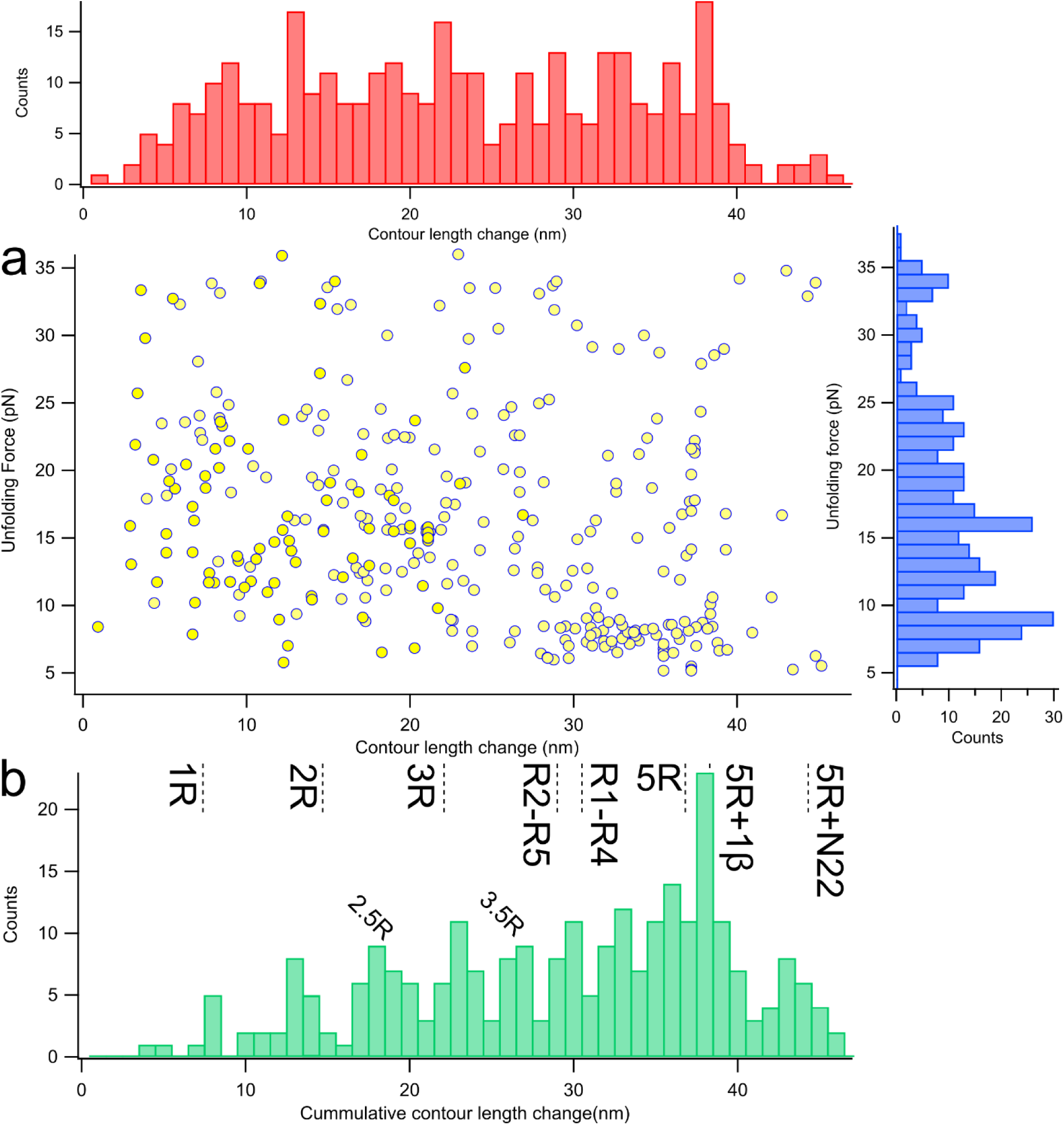
Analysis of discrete unfolding transitions. (**a**) Scatterplot of all observed unfolding forces and contour length changes and the corresponding force (right) and length (top) distributions. (**b**) Cumulative discrete contour length change (Δ*L_c__*_tot_) observed in each FEC. Labels indicate the amount of folded CsgA structure (measured from the AF3 model) consistent with the given Δ*L_c__*_tot_ value.

Entirely cooperative unfolding of the entire CsgA molecule (Δ*L_c_* around 44 nm in **Fig. 3a**) was rare, as was the complete unfolding via one or more discrete transitions (Δ*L_c_* around 44 nm in **Fig. 3b**). In most cases, the total contour length change, including both the shoulder lead-in followed by discrete transition(s) of 44 nm was consistent with that expected for the entire folded CsgA, spanning N22-R5, but with various unfolding pathways.

### Non-discrete transitions equilibrium model

FECs with no discrete transitions instead show numerous short, rapid oscillations within a narrow range of forces (few pN). Rather than fitting individual transitions, multiple FECs (with the same trap velocities, refolding times, and from the same molecule) were combined into ‘concatenated FECs’ for further analysis. Most refolding FECs also showed only a non-discrete transition and were similarly analyzed. The refolding curves largely retraced the unfolding ones (**Fig. 4a**), with some hysteresis evident. Thus, the unfolding and refolding traces were analyzed separately. To characterise the FECs showing only a non-discrete transition (shoulder), we used a model that describes sub-states in rapid equilibrium (**eqn 2**). We optimized the fitting to determine the minimum number of free parameters in the fitting function to adequately fit the data, finding that two transitions were optimal (a single transition severely underfit, while three transitions improved the fit negligibly, **Fig. S5**). The unfolding (**Fig. 4b**) and refolding (**Fig. 4c**) traces were both well-fitted as two transitions with nearly identical *F*_1/2_ and Δ*L_c_* values in either direction (**Fig. 4d**), supporting that these transitions occurred at near equilibrium. The lower-force (*F*_1/2_ = 4.3 ± 0.2 pN) transition corresponded to a larger conformational change (Δ*L_c_* = 27.7 ± 0.8 nm) compared to the higher-force (*F*_1/2_ = 7.2 ± 0.3 pN) transition (Δ*L_c_* = 13.9 ± 0.7 nm). Importantly, the average total Δ*L_c_* for these non-discrete transitions was 42 ± 1 nm, close to that expected (44 nm) for the entirely folded CsgA molecule. The pulling velocity had a small effect on the *F*_1/2_, with no significant impact on the Δ*L_c_* (**Fig. 4e**), consistent with these transitions occurring under rapid equilibrium conditions. Collectively, the low- and high-force transitions spanned a three-fold range of forces (3-9 pN). The free energies (*ΔG*) for the low- and high-force transitions, averaged across the unfolding and refolding curves (**Table S1**), were 4.1 ± 0.2 kcal/mol and 4.8 ± 0.3 kcal/mol, respectively.

**Figure 4.**
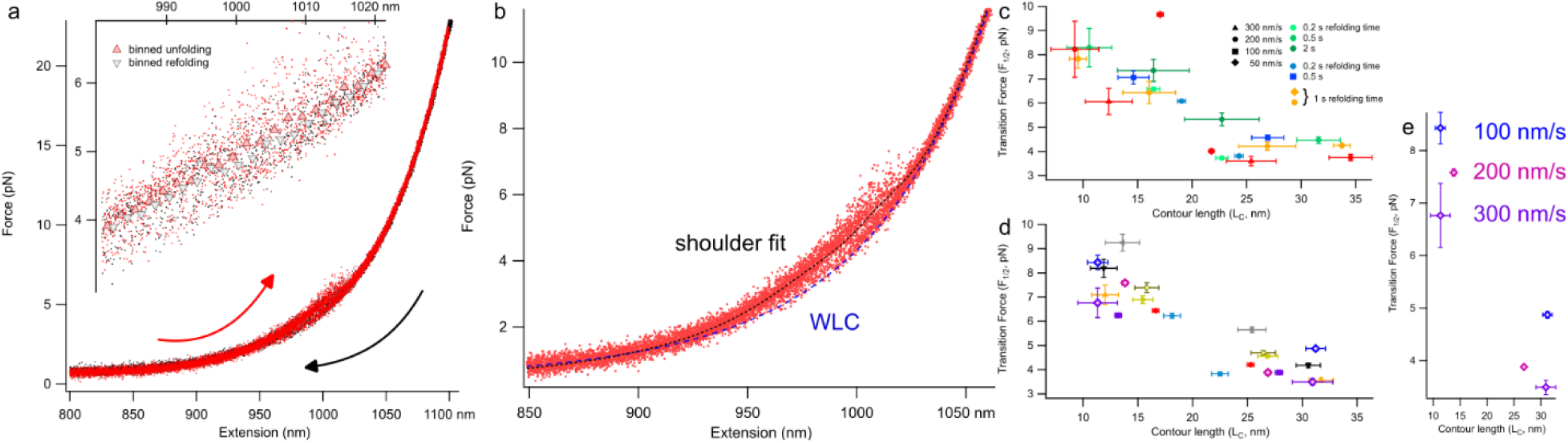
Non-discrete unfolding and refolding transitions. Multiple FECs were concatenated and fit with equation 2. (**a**) comparison of unfolding and refolding data revealed slight hysteresis, suggesting the shoulder feature represents near-equilibrium transitions. The unfolding (**b**) and refolding (not shown) data were both well fit assuming two independent transitions yielding *F*_1/2_ and Δ*L_c_* values for each transition. Fit results for (**c**) unfolding and (**d**) refolding, and the effect of pulling velocity on the transitions (**e**).

### CsgA refolding at constant trap separation

We also measured system-equilibrium refolding data with constant trap separation (CTS) over the course of five minutes (**Fig. 5**). For this experiment, FECs were first measured on a molecule, then with the molecule fully unfolded, the tension was gradually lowered to ~11 pN and the trap positions were held constant. CsgA was observed to refold under constant trap separation, reaching a final dynamic folding state between 14.4 and 15.3 pN, with contour length changes attributable to variable folding of the N22 domain onto R1-R5 (with all five R domains folded). The intermediate states observed are consistent with the lengths in terms of full and half integer multiples of R-domains. Near the third minute of the measurement, there was ~500 ms folding from 2R to 3.5R to 4R, before unfolding back to 3R domains, then down to a 2R domains steady state over a period of ~9 s. Also noteworthy is a ~6 s excursion to a fold of ~2.5R, up from 2R, before collapsing to all five R domains. Histograms of the endpoint folded region show three main states, with the 5R folded state fluctuating between an intermediate and the fully folded state (N22-R5) that, tentatively, we attribute to R1-R5 gaining parts of the N22 region dynamically, varying 5 nm in contour length change, or about 15 amino acid residues.

**Figure 5.**
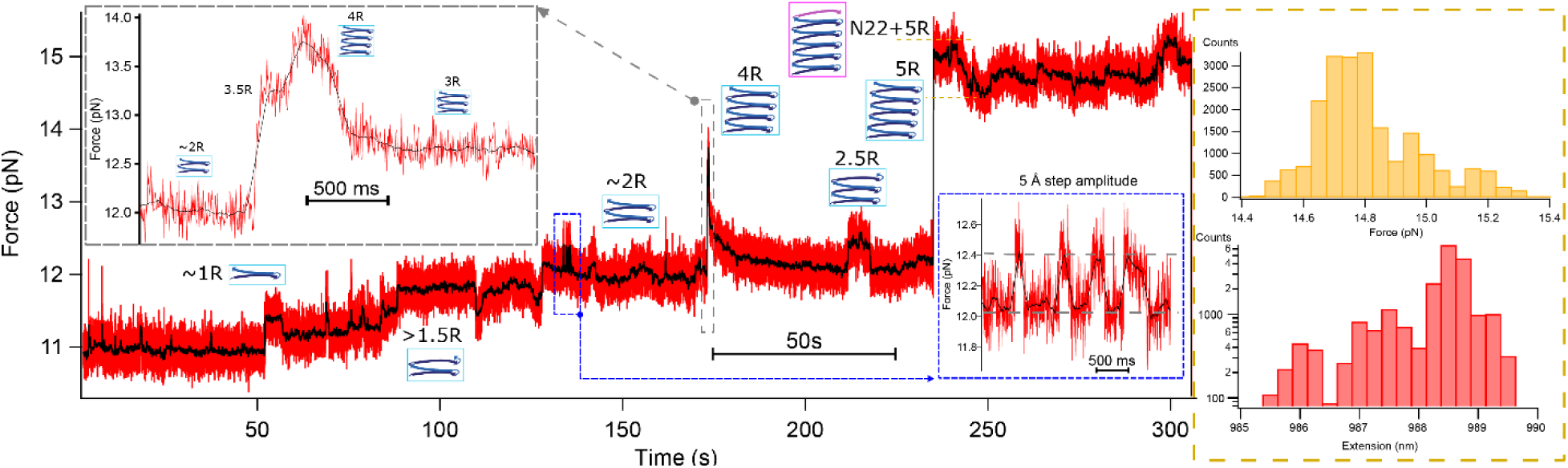
Equilibrium refolding of CsgA under tension. The force trace for a constant trap separation (CTS) experiment where each trap was stationary. CsgA was initially fully unfolded by force extension, then relaxed back to ~11 pN where trap positions were then fixed. With CTS, as the protein folds, force increases and extension decreases. CsgA folded up to a dynamic state from 14.4-15.3 pN (corresponding force and extension histograms for this region shown) before the tether broke. **Inset (left)**: a short (2.4 s) excursion from 2R to 3.5R up to 4R and finally to 3R. The 3R state sheared down to 2R, momentarily folded to 2.5R before folding into 5R+ (fully folded). **Inset (right)**: hopping between 2R and 2.5R.

### Steered MD simulation

Steered molecular dynamics simulations were carried out on the wildtype CsgA sequence with Cys linkers and the His-6 tag (**Fig. S8a**). The initial folded conformation was obtained using AF3^16^. The average work required to extend CsgA to the full 299 Å was 1313 kcal/mol, which is similar to previously reported values^40^. Protein contact maps (**Fig. S9**) indicate mostly sequential unfolding of the β-sheet secondary structures from the N-terminus. The R4 and R5 regions tend to form anti-parallel sheets before unfolding (**Fig. S9**). The force-extension curve averaged over all trajectories reveals five peaks that occur with a periodicity of approximately 75 Å. The peaks increase sharply and are followed by a slower decline, particularly for the unfolding of the last three regions (**Fig. S8b**). **Fig. 6a** shows a typical trajectory. In this trajectory, unfolding begins with the N-terminus and proceeds sequentially, where R4 and R5 restabilise into a misfolded anti-parallel sheet structure before complete unfolding.

**Figure 6.**
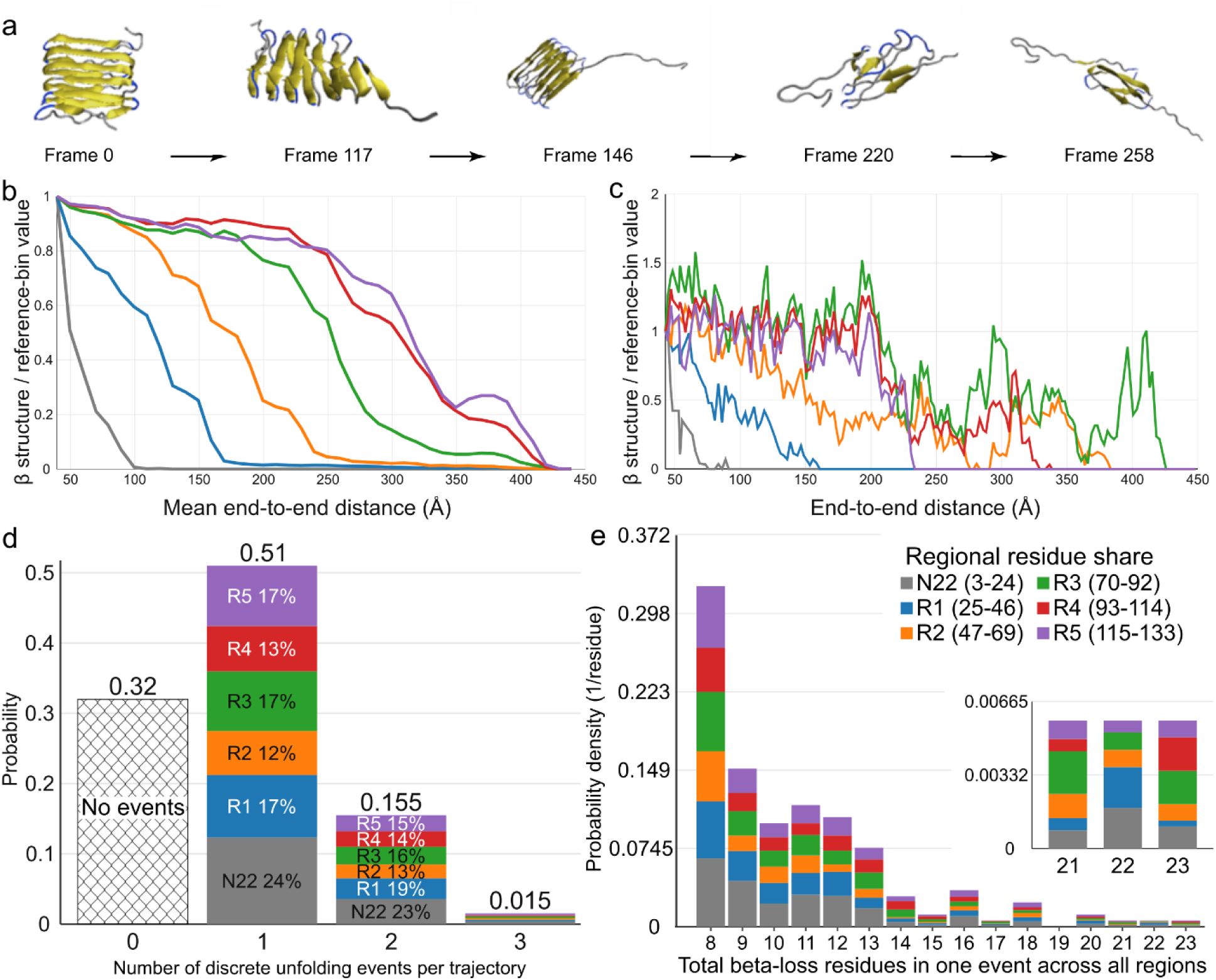
CsgA unfolding with steered molecular dynamics simulation. (**a**) Example structures from the unfolding trajectory. **(b)** Average probability of unfolding as a function of extension. (**c**) A single trajectory with discrete unfolding events. (**d**) Fraction of discrete unfolding events. (**e**) Cumulative frequency of discrete unfolding events. (**b, d, e**) Data averaged across 200 runs. Colouring in (**b-e**) according to the legend in (**e**).

Apart from this general trend, several unfolding and refolding patterns emerge, both continuous and discrete. In general, N22 unfolds first, while for regions R1-R4, unfolding begins at 0.5R intervals, i.e., when half of the previous region has unfolded (**Fig. 6b**). However, R5 unfolds together with R4 and restructures around 350 Å extension before complete unfolding. While the average unfolding trends appear relatively smooth, individual trajectories show discrete unfolding and refolding events (**Fig. 6c**). The number and frequency of such discrete events show strikingly similar trends to experimental data (**Fig. 6d**). The events appear to occur with relative uniformity across all regions. **Fig. 6e** shows the cumulative frequency of discrete unfolding events. As with the experimental results, the discrete events appear to have a periodicity, but on the order of about 4 residues (or 0.25 R). The simulations are dominated by single 8-residue events during the unfolding pathway, which occur at any one of the regions with nearly equal probability.

## DISCUSSION

The CsgA monomer was repeatedly stretched and relaxed using OT (**Fig.’s 2-4**). This data indicates that, as a monomer, CsgA exists in a dynamic equilibrium between a molten-globule collapsed state, a collapsed state with partially folded regions (1R to 4R), and the fully folded state spanning N22-R5 (**Fig. 7**). We speculate that these folded states rapidly aggregate and thus have evaded characterisation at the ensemble level. The CsgA monomer is generally viewed as being intrinsically disordered, adopting the beta-solenoid fold only upon incorporation into the curli fiber^11,18,41,42^. Our results alter this view, demonstrating that monomeric CsgA predominantly samples a range of metastable folded states, including a small fraction that are consistent with the fully structured β-solenoid fold. Our data are consistent, however, with the notion that CsgA folds into the β-solenoid structure with slow kinetics in the absence of a template or nucleation site^41^.

**Figure 7.**
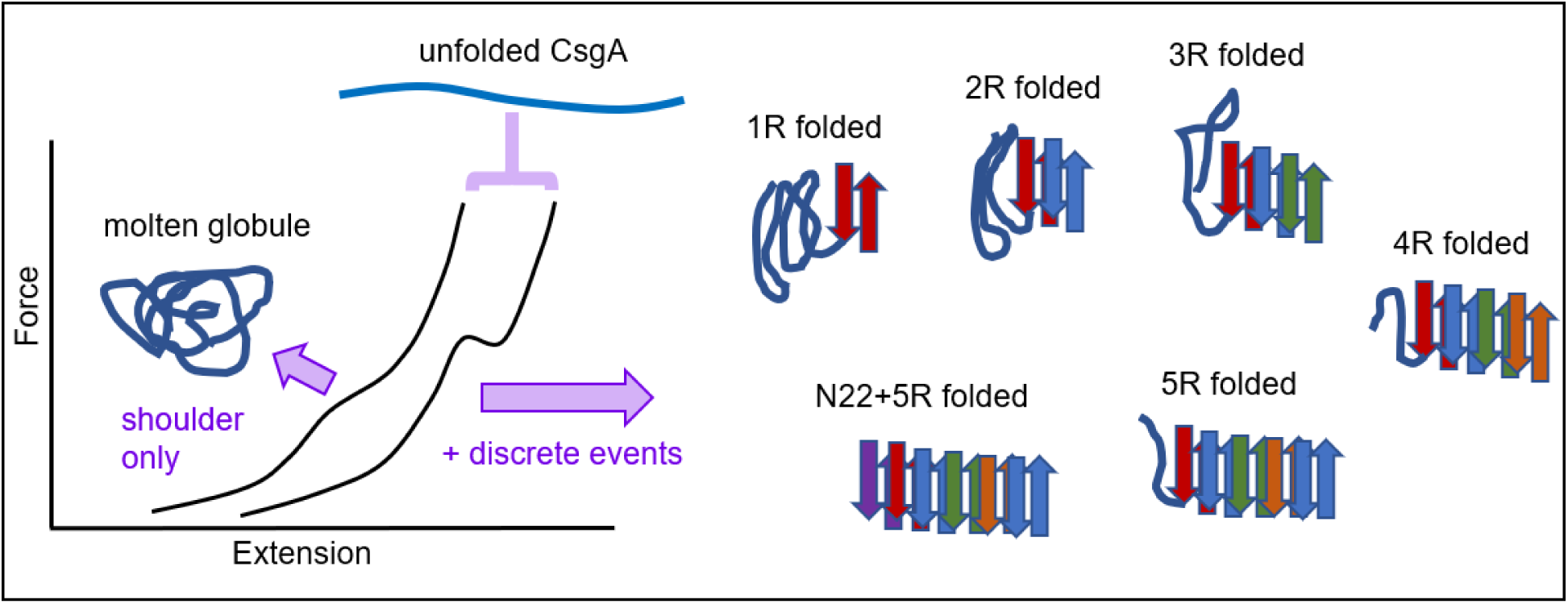
Schematic summary of CsgA folding dynamics. CsgA always shows some type of unfolding event: shoulder only, discrete transition, or shoulder + discrete transition. Δ*L_c_*__tot_ for the shoulder is consistent with the contour length change expected of a fully collapsed structure (e.g. involving all regions of CsgA). Taken together, this picture of single CsgA molecules sampling various metastable partially and completely folded conformations and collapsed yet amorphous (or ‘molten globule’) states is consistent with the long-held view of CsgA as an intrinsically disordered protein that comes from ensemble data. Our data, however, shows that CsgA can form a fully folded structure on its own, one that is consistent with the β-solenoid structure predicted for CsgA monomers. This structure is metastable in isolation and is likely stabilized in the oligomer form.

The always non-monotonic stretching behaviour of CsgA is markedly different from that of a typical IDP such as α-synuclein^26^, which mainly (~85% of FECs) showed featureless stretching FECs similar to that drawn in **Fig. 1d**. However, a shoulder feature seen in 30% of CsgA FECs was also observed in α-synuclein monomers (rarely) and tandem oligomers (predominantly)^34^, and also in tandem prion protein dimers^28^. We used a model that describes sub-states in rapid equilibrium to characterise the non-discrete transitions, as this was previously used to characterise the phenomenologically similar transitions in α-synuclein^34^ and prion protein dimer^28^. These transitions reach equilibrium much faster than the pulling speed, explaining the (near) absence of hysteresis or effect of pulling speed on the (un)folding forces (**Fig. 4**). The minimal model to fit the CsgA shoulder feature includes two transitions in series of marginal stability. CsgC was recently shown to promote the formation of disordered monomeric pre-fibrillar states, shifting the equilibrium away from amyloid formation^10^. CsgC transiently binds these monomeric CsgA species, and these may correspond to one or more of the structures observed here. Future studies of CsgC with CsgA using this system may reveal these transient binding partners.

Given the lack of dependence of the % of FECs showing discrete transitions (~70%) as a function of waiting time (**Fig. 2e**), we can infer that the partially- and fully-folded states are in equilibrium with the collapsed but rapidly fluctuating states represented by the shoulder transitions. Based on the equilibrium (70%/30%) the folded states are only marginally more stable than the molten globule form (*e.g*. 9.4 kcal/mol vs 8.9 kcal/mol). The metastable folded CsgA conformation would be further stabilized upon forming amyloid via interactions with neighboring units similarly as observed for other repeat proteins^43^. The mechanical lability of monomeric CsgA may play a functional role inside the cell, given that it is translocated through the CsgG pore complex out of the cell^11,44^, where a low mechanical stability can facilitate secretion^45^.

Constant trap-separation data (**Fig. 5**) show the sequential and complete folding of a CsgA monomer with end state fluctuations likely due to N22 transient structure. The N22 domain of CsgA is variably predicted as disordered, half- or fully-structured within the CsgA monomer^14^ (**Fig 1b**) and we find evidence for all three states in the folded ensemble (Δ*L_c_* values peaked around 36 nm, 39 nm, and 44 nm, **Fig. 3b**). A fully folded N22 is supported by our re-equilibrated AF3 model used as the starting point for SMD (**Fig. 6**), yet this region is believed to be excluded from the fibril core^18,41^. Together, this data suggests that N22 may transiently adopt folded structure but must unfold during amyloid assembly to expose R1. The SMD results support that N22 readily unfolds under load (**Fig. 6b**). Such transient structure might slow aggregation and could, in part, contribute to slow fibril assembly at the N-terminal end^46^. Further, the SMD results suggest (assuming microscopic reversibility^47^), that CsgA folding typically begins from R5, working back towards R1 and finally N22 (**Fig. 6b**). Initially, however, an early-stage misfolding can occur (**Fig. 6a**); such events would likely contribute to the rapid fluctuating shoulder transitions seen by OT (**Fig. 4**), as these misfolding events would need to reverse themselves for native folding to proceed.

Finally, while shearing is more likely to produce the cooperative, discrete transitions, unzipping (or “tearing”) results in sequential bond-breaking and gradual unfolding^48^. The slow, steady unfolding from 3R to 2R over 9 s (**Fig. 5**) is consistent with unzipping. Similar to the OT experiments, the SMD results show a combination of discrete and non-discrete unfolding events, with a predominant discrete step size of eight beta-sheet forming residues and periodicity of four residues (**Fig. 6e**), which is consistent with the size of shearing clusters from previous simulations^48^.

## METHODS

### Protein expression

The CsgA_CC_ construct (**Fig 1a**) with engineered internal Cys (A43C, V120C), flanked by ybbR tags (DSLEFIASKLA) and GGGGS spacers and a C-terminal His_6_ tag was expressed and purified as previously described with some modifications^18,35^. NEB 3016 ΔslyD cells harboring a pET11d vector encoding C-terminal His_6_-tagged CsgA were grown to OD_600_ ≳ 1 in LB broth containing 100 µg/mL ampicillin at 37°C. CsgA expression was induced with 0.5 mM IPTG for 2-3 h, then cells were harvested (8000 rpm, 10 min) and stored at −80°C. Cell pellets from 1 L of the induced cell culture were resuspended in 20 mL lysis buffer (8 M GdnHCl, 50 mM K_2_HPO_4_, pH 7.3) per gram of pellet, sonicated for a few min (20 s intervals), and shaken for 1 h at RT. The supernatant of the cell lysate was collected following centrifugation (4·10^4^ × g for 40 min), then mixed with an equal volume of equilibration buffer (50 mM Na_2_HPO_4_, 300 mM NaCl, 6 M GdnHCl, 10 mM imidazole, pH 7.4) and then purified using FPLC and a *HisPur* Cobalt column (Cytiva). Protein was eluted (6 M GdnHCl, 50 mM Na_2_HPO_4_, 300 mM NaCl, 150 mM imidazole, pH 7.4). Fractions containing CsgA were pooled together and stored at 4°C prior to analysis. All purification steps were conducted under 6 M GdnHCl to prevent premature fibrillation.

### Bioconjugate Chemistry

CsgA was labelled with dsDNA for OT experiments using the ybbR handle kit from Lumicks (Netherlands) and following the manufacturer’s instructions. Briefly, the protein was labelled on either side with ssDNA CoA-oligos of 36 nt, followed by annealing at 37°C for 15 minutes with complementary-overhang dsDNA handles of 1506 bp each, one labelled with biotin and the other with digoxigenin. The construct was diluted ~100 fold before being reacted with 2.08 μm anti-dig beads and 1.14 μm streptavidin beads (Spherotech).

### Single Molecule Force Spectroscopy

A Lumicks C-trap G2 (dual beam/trap optical tweezers) instrument equipped with a five-channel microfluidic flow cell was used. After forming a tether, at least a dozen force-extension curves (FECs) were collected for worm-like chain (WLC) fitting to ensure there were no obvious multiple tethers. Flow was applied to clear free beads from the flow cell before moving to the fourth channel of the microfluidic cell where the disulfide bridge of CsgA_CC_ was broken using ~100 mM mercaptoethylamine (MEA). After establishing the break of the internal disulfide bridge (**Fig. S2**), the dumbbell was moved back into a running buffer of PBS with an oxygen scavenging system as previously described^28^ with 50 U/mL glucose oxidase (GOx), 140 U/mL catalase and 0.01% D-glucose, as well as 10-20 mM of MEA to keep the protein’s cysteines reduced.

Bead position within the traps was recorded using back focal plane detection with position sensitive detectors and sampled at 78.125 kHz with an NI-4472 acquisition card (National Instruments) with 64× anti-aliasing (5 MS/s per channel). Data was imported into Igor Pro 9 (Wavemetrics), where it was down-sampled to millisecond time resolution with appropriate filtering for anti-aliasing. FECs were extracted and analysed by fitting with a series of WLCs for the dsDNA and protein as described previously^28^. In the WLC, force, *F* and change in extension, *Δx*, takes the functional form:

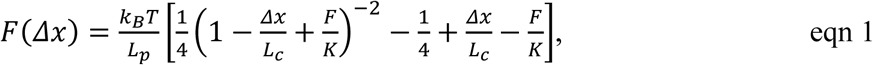

Where *k_B_* is the Boltzmann constant, with absolute temperature, *T*. The dsDNA fitting parameters have a persistence length (L_p_) of ~40 nm and elastic modulus (K) of ~700-1000 pN, while unfolded polypeptide lengths used an L_p_ of 0.85 nm, a K of 2000 pN and the crystallographic contour length of amino acids L_c_/a.a. = 0.36 nm, for conversion to involved residues^28^.

For the shoulder-like metastable features of the CsgA FECs, we have transitions that are both faster and on a similar timescale as data sampling and filtering of our instrumentation. We make use make use of the minimal model that assumes independent structures with two state transitions in rapid equilibrium, which has previously been successfully applied to α-synuclein^34^ and prion dimer^28^:

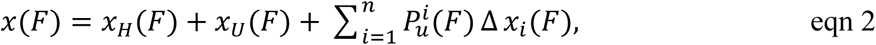

Where *x_H_*(*F*) is the extension of the dsDNA handles, *x_U_*(*F*) is the extension of the unstructured region of the protein, *n* is the number of two-state structures with distinct unfolding properties. For two-state unfolding at equilibrium, the probability term is described as:

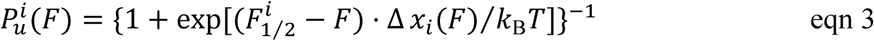

Where 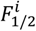 is the force at which the structure has a 50% probability of being unfolded, with the corresponding extension change for that structure Δ*x_i_*(*F*).

### Steered Molecular Dynamics

Simulations were performed with LAMMPS software^49^. The protein was modeled using the EEF1-SB implicit solvent force field^50^. CHARMM-GUI^51^ was used to add missing hydrogen atoms to the protein structure. VMD^52^ was used to visualise the system. Simulations were carried out in the NVT ensemble at 298.15 K with and integration timestep of 1 fs. The protein was placed at the center of a cubic simulation box of length 200 nm with periodic boundary conditions. Following initial equilibration, SMD simulations were carried out using a force constant of 50 kcal/mol/Å^2^ applied either to the N-terminus handle or the C-terminus handle with pulling velocity of 0.5 m/s. 100 independent SMD trajectories were generated for each case.

### Secondary-structure and beta-orientation classification

mkDSSP53 was used to classify the relevant secondary structures along the trajectory. Visual Molecular Dynamics (VMD) was used to colour each residue according to its mkDSSP assignment. Positional data from the LAMMPS trajectories were then used to classify each identified β-sheet segment with a minimum of two residues as parallel, antiparallel, or mixed. Potential sheet partners were separated by at least four residues and were required to contain at least one intersegment C-alpha-C-alpha distance of 6.5 angstroms or less. The dot product, *d*, between positional segment vectors was used to determine relative orientation. Segments were classified as parallel (*d* > 0.25), antiparallel when (*d* < −0.25), and mixed when a segment showed a parallel assignment with one partner and an antiparallel assignment with another.

### Discrete unfolding event identification

Discrete unfolding events were identified using residue-level β secondary-structure assignments. For each residue in a given frame, average β occupancy was evaluated over the preceding 10 frames and over the following 24-frames. The residue was considered persistently unfolded when its β occupancy was at least 50% before the given timeframe and less than 20% afterwards. The single residue unfolding transition was then confirmed over four additional frames, during which the residue was required to be non-beta in at least 75% of these frames. Finally, for an event to be considered part of a discrete unfolding event, at least eight residues underwent sustained beta loss within an eight-frame rolling window within the given 34-frame time interval.

### Regional folded-fraction analysis

Regional folded fractions were calculated by comparing the number of β-assigned residues in each frame with the corresponding count in the first frame. A residue was counted as unfolded only when it remained non-beta for at least 10 consecutive frames.

## Acknowledgements

We thank Matthew Chapman for sharing the *E. coli* strain used for protein expression, E. Debenedictis and S. Keten for sharing their computational models of CsgA structure, and Johannes Stigler for sharing the C-trap HDF5 import code for Igor. We thank Noel Q. Hoffer and Krishna P. Neupane for providing helpful discussions about optimising the optical tweezers experiments. We thank Fan Bu for protein expression protocol development, as well as Hrishika Dekate and Rachel Rosenberg for preparing protein samples. The UBC Bioimaging facility (RRID: SCR_021304) was used for TEM imaging.

## Data Availability

All data will be made available upon reasonable request.

## SUPPORTING INFORMATION

**Figure S1.**
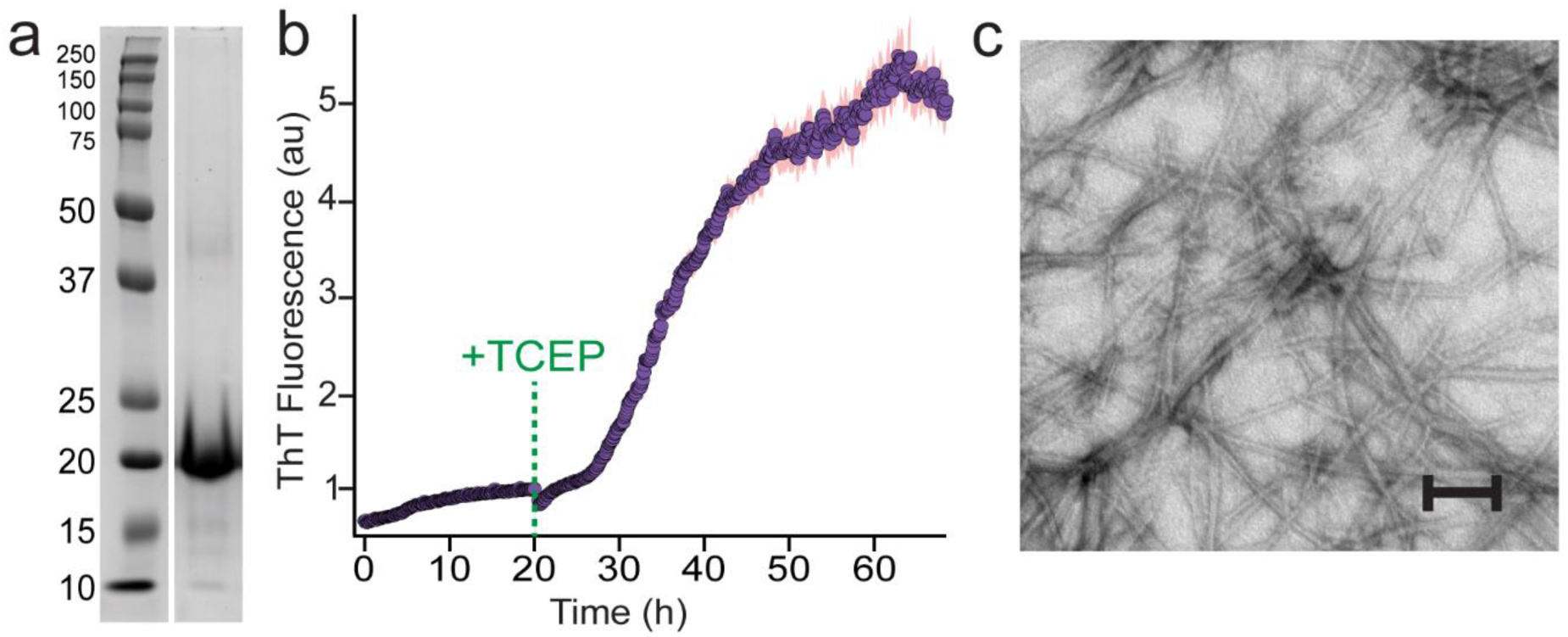
Ensemble characterisation of the yybR-CsgA_CC_-ybbR construct. **(a)** Reducing SDS-PAGE of the IMAC-purified protein. **(b)** The CsgA_CC_ mutant is kinetically trapped from aggregation by an intramolecular disulfide bridge. Amyloid formation was triggered by adding reducing agent and measured using ThT fluorescence. (**c**) TEM microscopy showed that the reduced construct formed amyloid fibrils. Scale bar = 100 nm.

### ThT aggregation kinetics

Zeba desalting columns (Thermo Scientific) were used to quickly buffer exchange the CsgA protein from 6 M GdnHCl into 50 mM HEPES buffer at pH 7.0. Thioflavin T (ThT) was added to a concentration of 20 μM before the sample was placed into a 96 well plate for bottom reads with a SpectraMax Gemini EM (Molecular Devices) with excitation at 450 nm and emission at 495 nm, 15 reads per cell, and five replicates without shaking. Tris(2-carboxyethyl) phosphine (TCEP), used to reduce the internal disulfide, was made into a pH 7 stock in HEPES.

### Optical trapping and reduction of the disulfide-linked CsgA

During screening, most tethers in the absence of reducing agent showed no unfolding events during pulling (**Fig. S2a**). A reducing agent was introduced by moving the dumbbells into a flow channel containing mercaptoethylamine (MEA), at which point when the cysteines became reduced, the contour length of flexible polypeptide then increased significantly, producing a notable elongation in the FEC, and then subsequent unfolding FECs may show discrete evens characteristic of cooperative R domain unfolding (**Fig. S2b**). However, before moving to the reducing buffer, CsgA would occasionally show a small unfolding event (**Fig S2c**) consisting of an unzip of 10 to 13 nm in the ~5 to 7 pN range. Outside of the disulfide bridge at 43C and 120C, R1 and R5 have 32 free amino acid residues, which normally would be part of the β-hairpin structure of their respective R domain. Residues 44 through 119 are bound into a loop and do not contribute to the contour length. Balistreri *et al.*^1^ hypothesized that the disulfide bridge keeps the CsgA in a disordered state, but the rapid onset of fibrillation of CsgA A43C/V120C after the introduction of a reducing agent (TCEP) suggests to us that some seeding aggregates may already be possible. OT results indicate, typically on a first pull alone, that a partial β-hairpin or β-sheet formation sequestering these 32 free residues is possible given the change in contour length. AF3 and RoseTTAFold do not offer a structure prediction with the disulfide intact. AF3 uses multiple sequence alignment on primary input, which will classify the R-domains without considering the A43C/V120C amino acid changes. We observe no indications of a structural contribution from the ybbR flanking sequences, which is consistent with the experiments of Synakewicz *et al.,* comparing cysteine and ybbR/Sfp attachment chemistry for effects on forces observed in FECs^2^.

**Figure S2.**
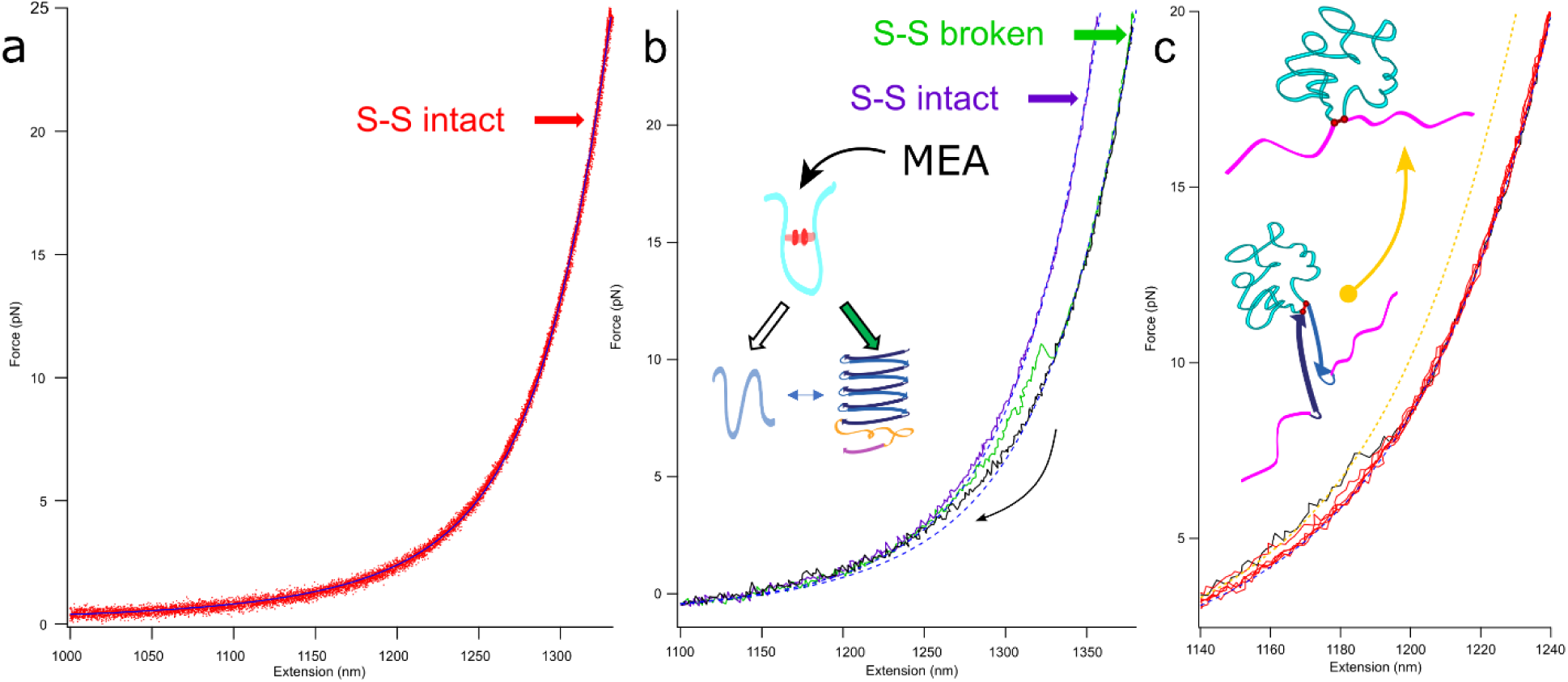
CsgA engineered for force spectroscopy studies. Optical tweezer data were fit with a WLC model to screen for single tethers. (**a**) Without reducing agent (violet; overlay of 14 FECs). (**b**) Moving the sample into the channel containing 100 mM MEA reduces the disulfide permitting complete unfolding (green) and refolding (black). (**c**) Structure formation within disulfide-linked CsgA: occasionally a 10-13 nm unzip is seen before the introduction of reducing agent, which is consistent with the free regions of R1 and R5 forming a folded structure outside of the disulfide looped region containing R2-R4. Dashed lines in all panels indicate WLC fits.

**Figure S3.**
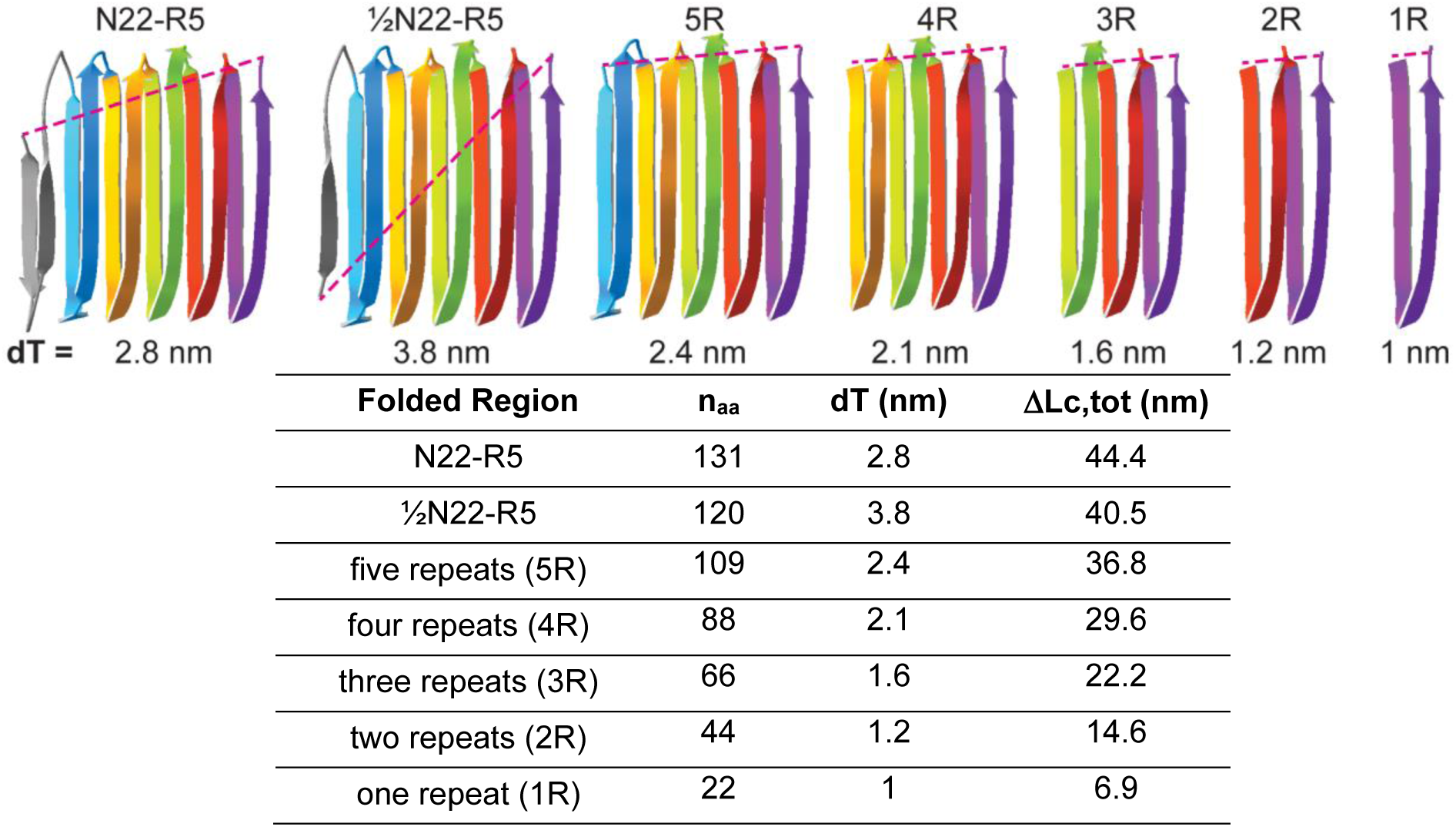
Example calculations of dT and ΔLc values from various CsgA folded regions. AF3 model used to estimate dT for different amounts of folded CsgA. The number of residues (n_aa_) and the distance between termini (dT) of the structure that folds/unfolds, along with the contour length per aa (0.36 nm) are used to calculate the contour length change for that structure (equation 1). The average n_aa_ was used in the cases of 4R through 1R.

**Figure S4.**
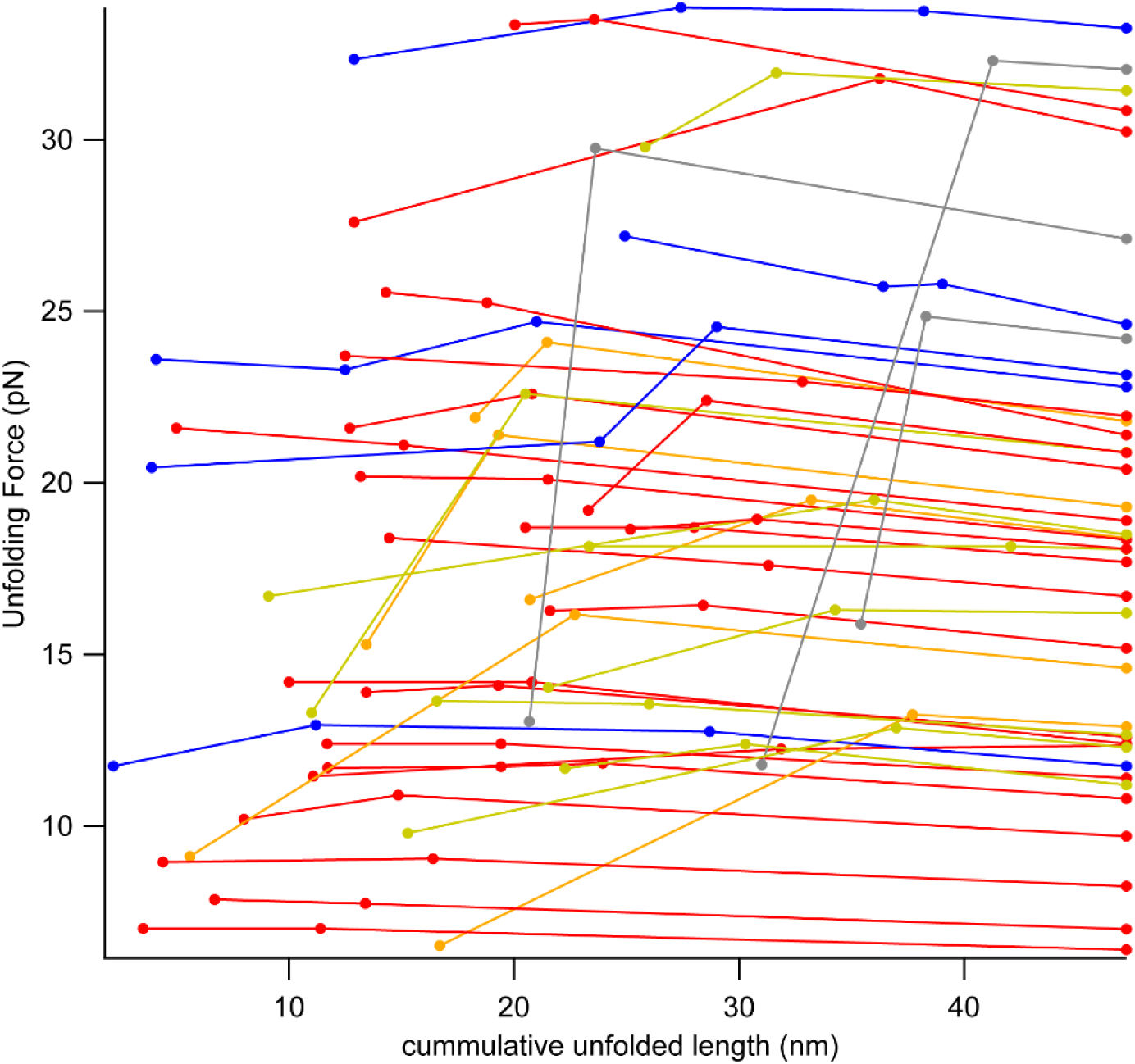
Discrete unfolding paths of CsgA. Sequential unfolding forces and cumulative contour length changes within individual FECs. In FECs tabulated from five molecules, multiple discrete unfolding steps can be observed (three distinct states in blue), up to the point where they unfold to 44 nm, which is the fully unfolded length of the CsgA construct. (Much of the contour length change from the fully folded state may not occur discretely, and thus not shown here.)

**Figure S5.**
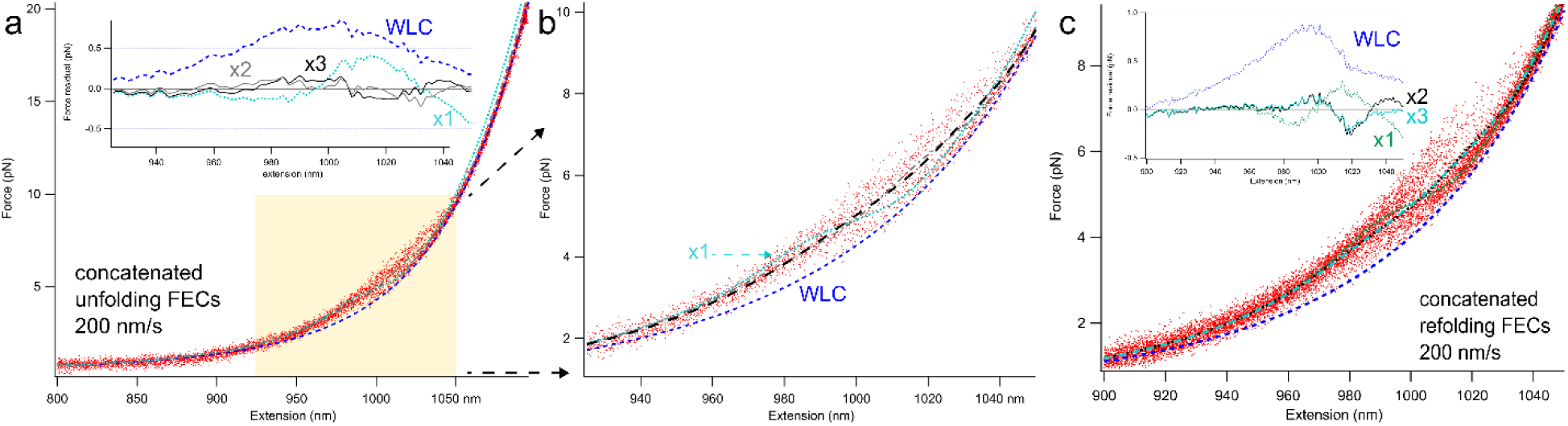
Fitting non-discrete transitions with an equilibrium model. Multiple FECs were concatenated and fit with equation 2. Both (**a**) unfolding and (**c**) refolding data were adequately fit assuming two independent transitions. Using only a single transition yielded a poor fit, while including three transitions did not improve the fit substantially, as judged by the fit residuals (top). (**b**) Expanded view of the refolding shoulder transition.

**Figure S6.**
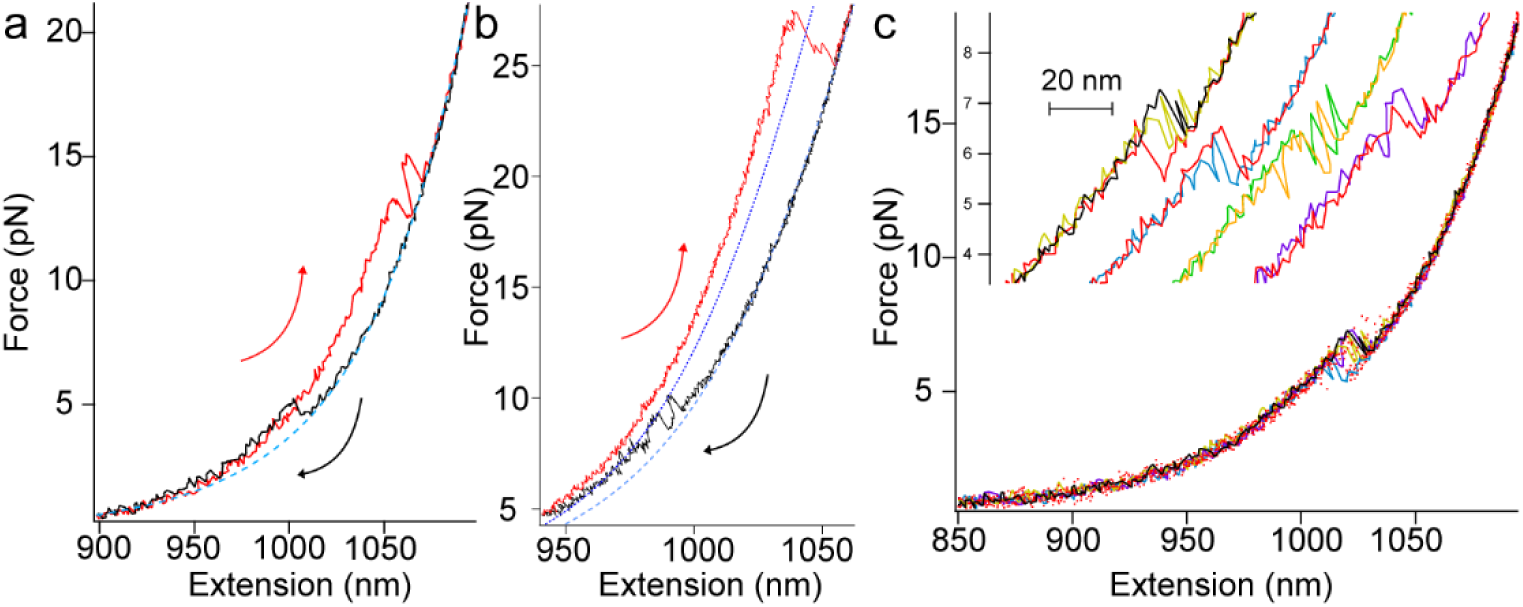
State switching and discrete refolding events. **(a)** Example FEC (red) of molecule that unfolded at 13.9 pN yielding 20.5 nm to the fully unfolded state, then refolded and unfolded again at 15.6 pN, yielding 18.6 nm. Refolding (black) and WLC for the unfolded state (light blue). **(b, c)** Discrete refolding events. **(b)** Discrete refolding contour length change of 15.6 nm can be observed to middle dotted line (dark blue), just over of the length of 2R domains occurring just below 10 pN. The unfolding curve in red, unfolds from 3R to ~1.5R to ~1R to fully unfolded near 25 pN. (**c**) Examples of discrete refolding events.

**Fig S7.**
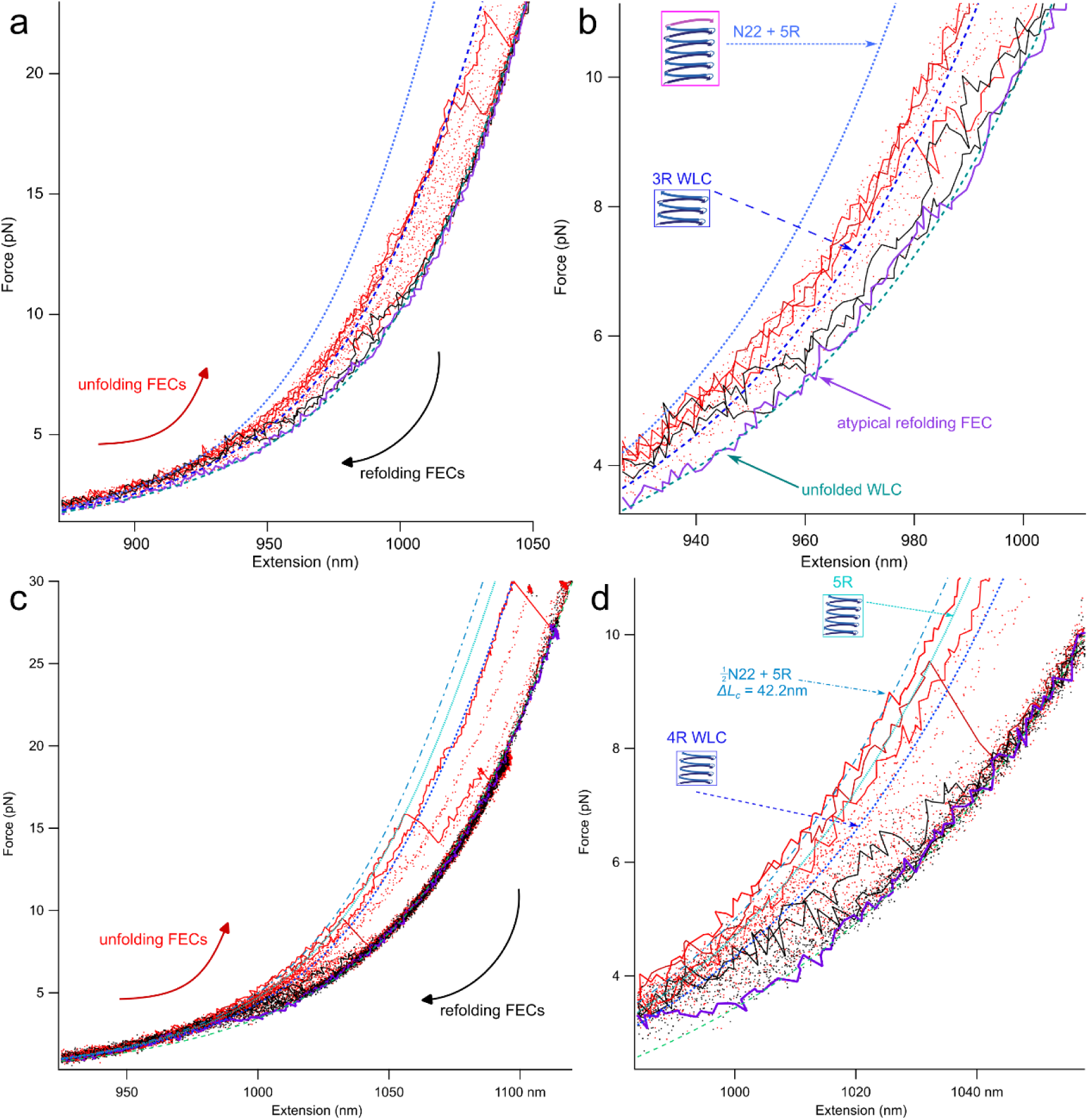
Fitting to the shoulder limits. (**a,b**) Aligned set of unfolding FECs (red) with two typical refolding FECs (black), displaying the shoulder behaviour, and an outlier featureless, monotonic refolding FEC (violet). Based on the refolding curves, the WLC for a 5R and N22 fully folded is overlaid in light-blue. (**b**) Zoomed view for clarity, the middle-WLC fit state has a *ΔL_c_* of 21.2 ± 0.2 nm, close to the 3R average. A gradient aligning to 5R or more of condensed *L_c_* until a discrete unfolding is very typical of our observed unfolding FECs. This is repeated with another molecule (**c,d**) with sufficient data and an outlier refolding FEC to fit from (violet). Here three folded states with discrete unfolding events are fit and shown, with lines for 4R, 5R and partial N22 (*ΔL_c_* of 42.2 ± 0.3 nm) shown in (**d**).

**Figure S8.**
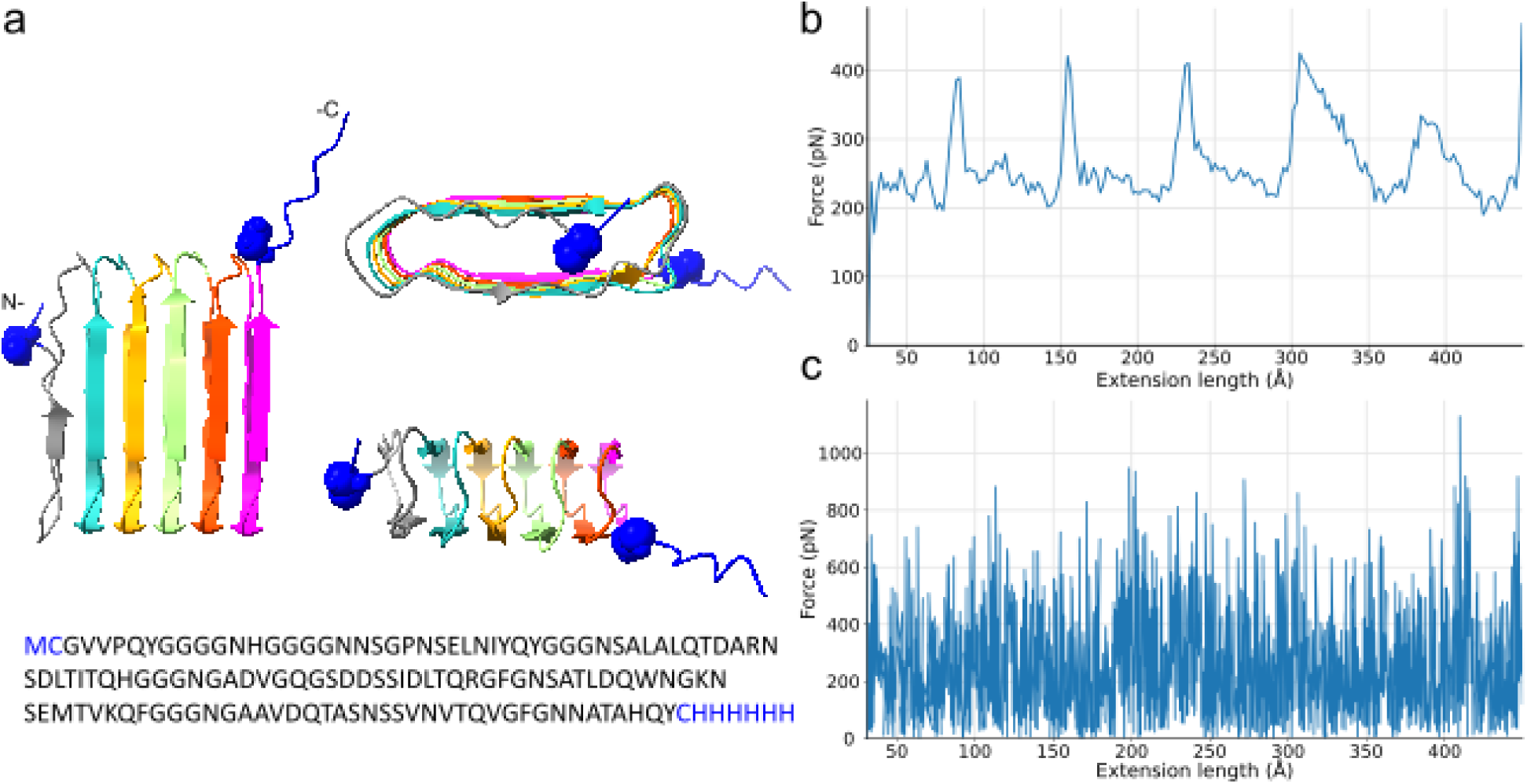
Steered molecular dynamics of CsgA unfolding. (**a**) CsgA construct. The mature CsgA wild-type sequence (black) was flanked by Cys (side chains shown) and a His-6 tag (blue). (**b**) Cumulative force extension curve averaged across 200 runs. (**c**) Force applied in a single run.

**Figure S9.**
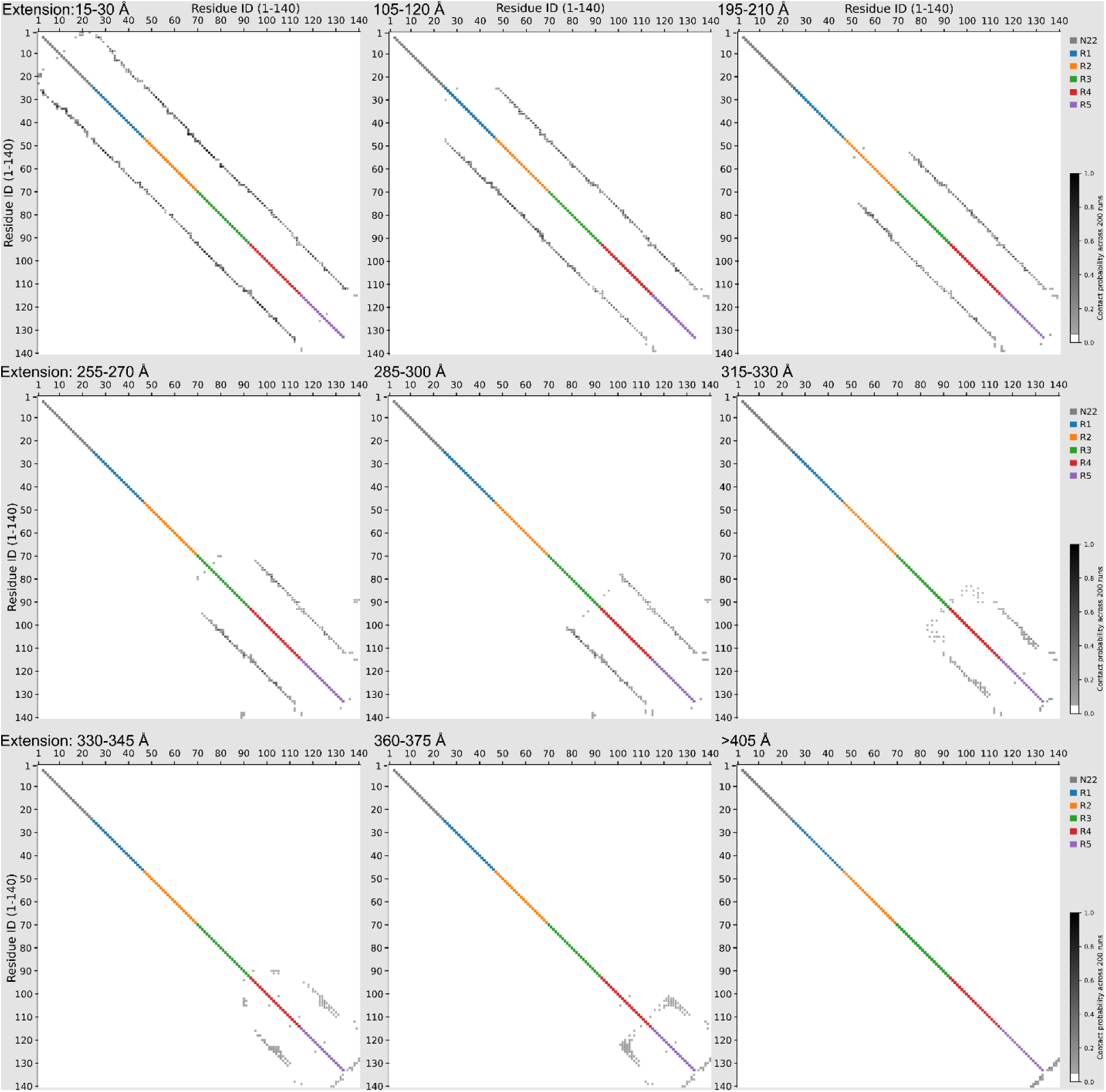
Protein Contact Maps (PCMs) from steered molecular dynamics simulations. Diagonal line coloured according to domain (N22, R1, R2, R3, R4, R5).

### Fitting the non-discrete transitions

The equilibrium free energies and model parameters were calculated for each molecule, both refolding (8) and unfolding (3) separately, then averaged over molecules. Measurements made by Synakewicz *et al*. showed a relative insensitivity to unfolding forces with respect to ramping rates from 10 to 1000 nm/s^2^. While they do observe a lack of hysteresis ≤ 100 nm/s, it becomes significant near and above trap velocities of 1000 nm/s. In our case, the majority of the data was sampled at 200 nm/s with some smaller sets at 50, 100 and 300 nm/s, these were averaged within the single molecule dataset itself. Uncertainties provided are the standard error of the mean for the results shown in the table below. Unfolding and refolding quantities overlap within error, indicating that CsgA’s quantifiable states are close to, if not at equilibrium within our ramping rates probed here.

The free energies of the structures, Δ*G* were determined from the values for *F*_1/2_ and Δ*x*(*F*_1/2_), where Δ*x*(*F*_1/2_) was obtained from the WLC fit at *F*_1/2_ for a protein of length Δ*L_c_* from the fit results of eqns 2 and 3. From here the structure energy, Δ*G* comes from: Δ*G* = *F*_1/2_·Δ*x*(*F*_1/2_) − [Δ*G*(*x*)]_stretch_, where [Δ*G*(*x*)]_stretch_ is the energy required to stretch unfolded protein to *F*_1/2_^3^. Given that [Δ*G*(*x*)]_stretch_ is defined as 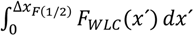, we can calculate Δ*G* from the inverse: 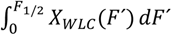, from Young’s inequality of integration. Energies were calculated for a temperature of 24 °C.

**Table S1.**

| substate | $\Delta L_c$, nm | $F_{1/2}$, pN | $x(F_{1/2})$, nm | $\Delta G$, pN·nm | $\Delta G$, kcal/mol |
| --- | --- | --- | --- | --- | --- |
| Unfolding FEC |  |  |  |  |  |
| small | $13 \pm 1$ | $7.17 \pm 0.10$ | $7.1 \pm 0.6$ | $31 \pm 3$ | $4.5 \pm 0.4$ |
| large | $28.1 \pm 0.9$ | $4.39 \pm 0.05$ | $11.8 \pm 0.3$ | $30.0 \pm 0.5$ | $4.32 \pm 0.07$ |
| Refolding FEC |  |  |  |  |  |
| small | $14.1 \pm 0.8$ | $7.3 \pm 0.3$ | $7.7 \pm 0.9$ | $34 \pm 2$ | $5.0 \pm 0.3$ |
| large | $27.6 \pm 1.1$ | $4.2 \pm 0.2$ | $11.3 \pm 0.4$ | $28 \pm 2$ | $4.0 \pm 0.3$ |

